# From dust to seed: a lunar chickpea story

**DOI:** 10.1101/2024.01.18.576311

**Authors:** Jessica A. Atkin, Sara Oliveira Santos

## Abstract

Food sustainability is one of the most significant barriers to long-term space travel. Providing resources from Earth is not cost-efficient, and resupply missions are not viable to meet the needs of long-term life in deep space conditions. Plants in space can provide a source of nutrition and oxygen, reducing the reliance on packaged foods, reducing resupply needs, and extending the duration of missions. Using lunar regolith simulant, we employ a novel methodology to create a sustainable and productive growth medium to support the cultivation of horticultural crops on the Moon. Implementing microbial soil regeneration mechanisms derived from Earth, we leverage the interaction between Arbuscular Mycorrhizal Fungi (AMF) and Vermicompost (VC) to create a fertile LRS matrix. These amendments can sequester toxic contaminants, improve soil structure, and increase plant stress tolerance. We demonstrate the ability to produce chickpea (Cicer arietinum) in lunar regolith simulant augmented with AMF and VC under climate-controlled conditions. We cultivated chickpea to seed in a mixture containing 75% Lunar Regolith Simulant. Preliminary results suggest that higher LRS contents induce heightened stress responses. However, plants grown in 100% LRS inoculated with arbuscular mycorrhizal fungi demonstrated an average two-week survival extension compared to non-inoculated plants. This study provides, for the first time, a baseline for chickpea germination in varying mixtures of LRS and VC and will inform future studies as humanity goes back to the Moon.

## Introduction

To return humans to the moon and establish a lunar presence, we must maximize in-situ resources and use lunar regolith (LR) and regenerative processes to provide a sustainable support substrate for horticultural [1]. Although LR shows traces of nutrients required for plant growth in plant-inaccessible forms, the use of LR as a sole growing medium presents challenges due to toxic elements, contamination from radiation, lack of organic materials, an absence of rhizosphere microorganisms, and poor chemical and physical properties [2–4].

Experiments using LR samples returned from the Apollo missions have investigated plant growth in the presence of lunar materials. Studies showed that during short-term exposure to irradiated lunar samples, seeds could successfully germinate [5, 6]. Research using *Arabidopsis thaliana* showed that although plants can germinate in LR, they exhibit slower development and severe stress morphologies [7]. Long-term exposure to metals such as those in LR will contribute to plant toxicity, severely altering physiological processes and causing oxidative damage. Furthermore, lunar regolith has poor physical and structural properties. Water permeability of lunar highland regolith is one to two orders of magnitude lower than that of fine silica sands and can be attributed to the irregular and angular shapes of LR due to lack of weathering [8]. These properties, particle size (0.25 mm to 1 mm), and poor aggregate development pose challenges to water retention, nutrient availability, and gas exchange [9]. Additionally, the Lunar regolith lacks a soil microbiome crucial for nutrient transformation and plant uptake, preventing efficient nutrient conversion and plant availability. These combined factors highlight the limitations of cultivating plants using a lunar-based substrate.

To use LR as a growth medium, we must first initiate an LRS matrix transformation to enhance soil structure and mitigate element toxicity. This involves incorporating stable organic amendments to improve the structural properties of the regolith and introduce a microbiome. Studies using mixtures of compost and regolith simulant showed that amendments can improve Martian and Lunar regoliths as a plant growth medium by improving nutrient availability and hydraulic properties [10–12]. Studies using both Martian and Lunar regolith, in comparison to Earth soils, show that germination rates are highest in Martian soil simulant and lowest in Lunar regolith simulant, which may be attributed to the larger water holding capacity of Martian soil simulant in comparison to LRS [13]. Due to its physical properties, lunar regolith has proved to be a challenging growth medium, even with compost mixtures. To succeed in using LR as a growth medium, we must change its chemical and physical properties to facilitate the establishment of a microbiome. Here, we focus on overcoming these challenges by creating a fertile support matrix to sustain plant and microbial life by using arbuscular mycorrhizal fungi (AMF) and vermicompost (VC).

AMF can remediate heavy metal-contaminated soils through phytoremediation and safeguard the plant through multiple mechanisms. Exudates from extraradical mycelia are used to sequester heavy metals (HMs) in the rhizosphere, reducing their bioavailability and preventing them from uptake by the host plant. In conjunction, extracellular polymeric surfaces on the hyphae surface work with other compounds to prevent the entry of HMs into the root tip (Figure 1) [14, 15]. A secondary mechanism allows AMF to accumulate heavy metals in their biomass and the host plant’s roots [16]. This enables them to stimulate plant resistance, reduce the negative impact of heavy metal toxicity, and promote plant growth under conditions of metal stress [17]. AMF plays a vital role in regulating rhizosphere productivity by enhancing nutrient cycling capabilities and promoting soil aggregation, thereby influencing particle organization and structure [18, 19]. Moreover, AMF produces glomalin, a glycoprotein that acts as a binding agent, improving aggregate stability and reducing soil erosion [20].

**Figure 1:**
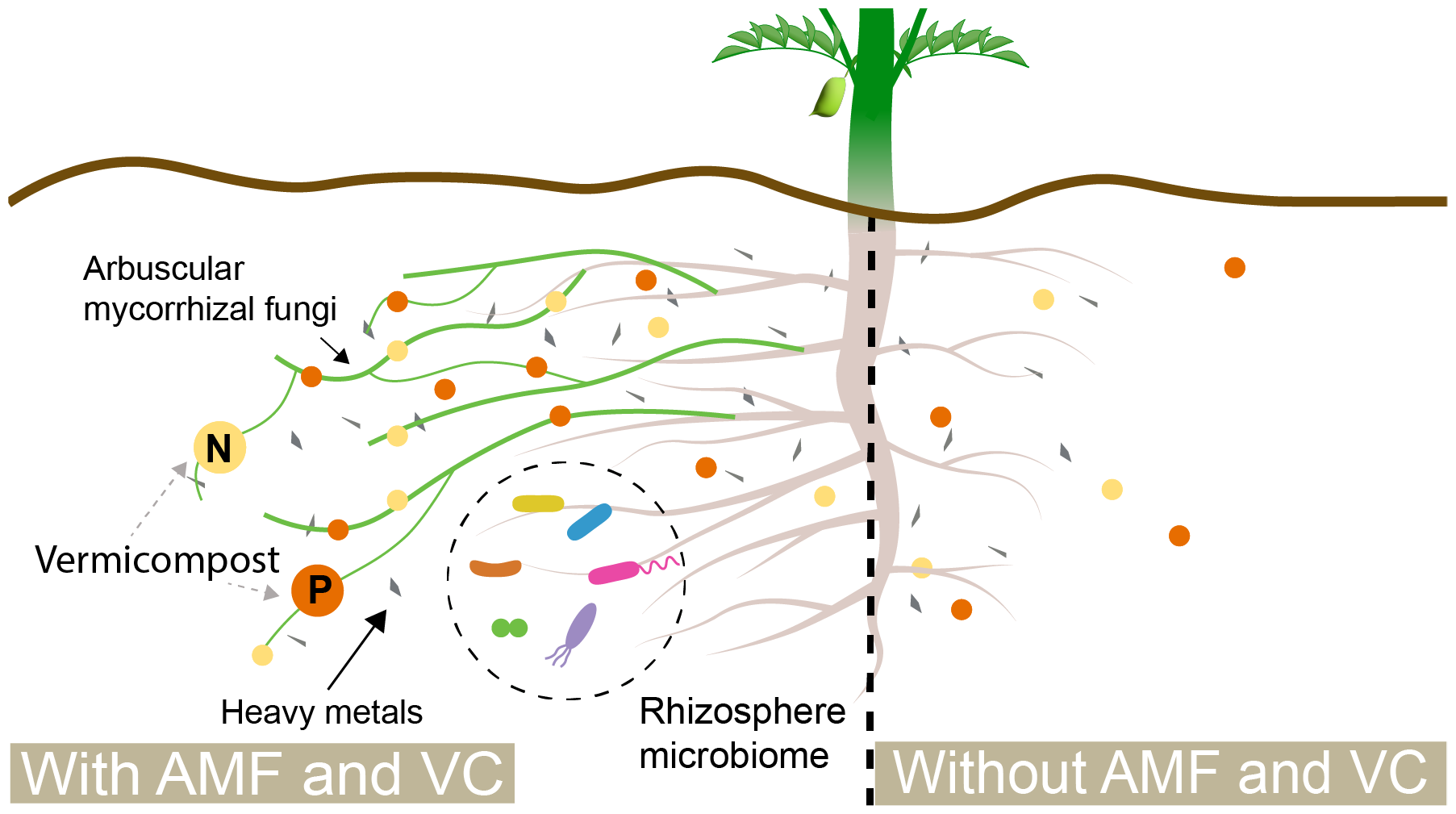
*Cicer arietinum*, AMF and VC in the rhizosphere. Chickpea has a taproot reaching up to 30 cm with lateral root branching. The root with AMF has an extended surface area reaching otherwise inaccessible nutrients and sequestering heavy metals (HMs) from uptake. The root structure without AMF has less surface area, shorter roots, and fewer protective mechanisms against HMs.

*Vermicomposting* is a bio-oxidative process resulting from the synergistic action of red wiggler earthworms (*Eisenia fetida*) and microorganisms to decompose biowaste. The resulting biostimulant is rich in plant essential nutrients and minerals and has a diverse microbiome. Earthworms produce water-soluble nutrients and humus while improving soil aggregate formation, lowering bulk density, modifying soil structure, and improving water retention. [21]. Vermicompost is an effective way to recycle clothing, hygiene items, and food wastes found in NASA’s logistics reduction and repurposing project to produce a sustainable, microbe-rich fertilizer.

We use chickpea (CP) as our model plant due to its nutritional content and ability to host symbiosis with AMF [22]. CP is a legume crop high in protein, carbohydrates, iron, phosphorus, calcium, vitamin B, and other nutrients and does not require large water or nitrogen inputs. CP is utilized globally as a nutritious protein substitute for meat, and has been used widely in studies evaluating the remediation of radioactive and metal-contaminated soils[23].

We hypothesize that the symbiosis created through AMF, VC, and CP will provide a sustainable support substrate for lunar crop production. We test this hypothesis through experiments using AMF in varying concentrations of LRS/VC.

## Methods

Experiments were conducted over 120 days in a mylar-lined grow tent with controlled temperature (≈23*°* C) using full spectrum adjustable lighting (4000W LED Grow Light Full Spectrum) employing a 17-hour photoperiod (380-730nm), and using a complete block design (Figure 2). Given the limited availability of Lunar Regolith obtained from the Apollo missions, Lunar Regolith Simulants (LRS) have been engineered from Earth’s geological materials. The simulants replicate specific Lunar landing site characteristics by incorporating plagioclase, pyroxene, olivine, ilmenite, chromite, and glass. Here, we use a high-fidelity lunar regolith simulant, LHS-1 (ExolithLabs, Oviedo, FL), derived from the lunar highlands [24]. This LRS replicates the mineralogy, geochemistry, and particle size of LR [25–27]

**Figure 2:**
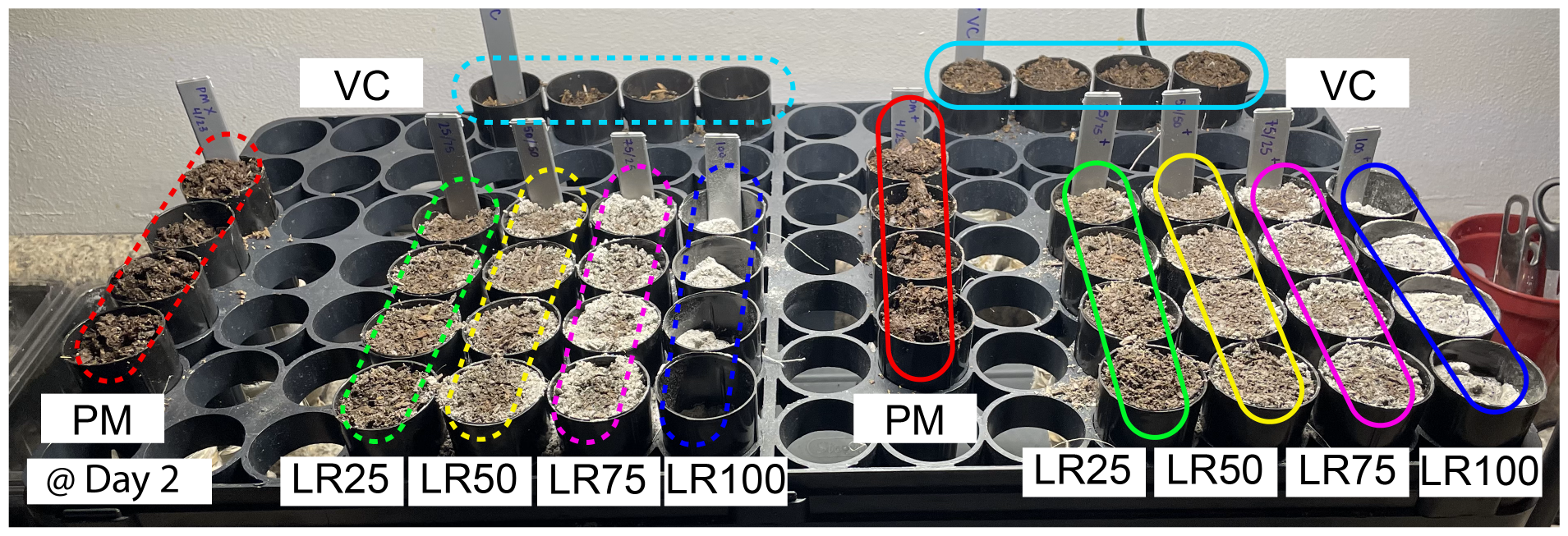
Experimental setup using a complete block design, with one seed planted in each container. Two controls were used: potting mix (PM) and vermicompost (VC). Four different compositions of Lunar regolith simulant and vermicompost were used: LR25 uses 25% LRS/ 75% VC, LR50 uses 50% LRS/ 50% VC, LR75 uses 75% LRS/ 25% VC and LR100 100% LRS. The dashed line shows seeds that have not been inoculated with Arbuscular Mycorrhizal Fungi, and the full line shows seeds that have been inoculated.

Plants were grown in 20 cm tall containers with a total volume of 107 *cm*^3^ (Ray Leach Cone-tainer) to allow for root expansion over their growth period (100 days) to full maturity. The Desi chickpea variety was chosen for its compact size and tolerance to stress [28]. Cheesecloth was affixed to the lower opening of the containers to prevent the loss of small particles. Two controls were used, one using potting mix (PM) (n = 8) and one using vermicompost (VC) (n = 8) (Black Diamond Vermicompost, Paso Robles, CA). VC and LRS were mixed by weight (*ρ*_*V C*_ = 0.39 *g* /*cm*^3^, *ρ*_*LRS*_ = 1.3 *g* /*cm*^3^) in four different compositions: LR25, 25% LRS/ 75% VC (n = 8); LR50, 50% LRS/ 50% VC (n = 8); LR75, 75% LRS/ 25% VC (n = 8); and LR100, 100% LRS (n = 8) (Figure 2). Half of the compositions (n = 24) were inoculated with Arbuscular Mycorrhizal Fungi (Mycoapply, Grants Pass, OR) containing *Rhizophagus intraradices, Funneliformis mosseae, Claroideoglomus claroideum* and *Claroideoglomus etunicatum*. The granular inoculum was placed in direct contact with the seed as a coating. Inoculations were performed at twice the label rate to account for limitations in colonization due to LRS toxicity (per the manufacturer’s recommendation). Seeds were planted in an LRS matrix and evaluated 21 days after 50 % of seedling emergence in the control groups [29]. At week 10, the light spectrum (purple, 380, and 450 nm) was adjusted to induce reproductive growth.

## Results and discussion

All seeds were planted on day 0 and watered with 4 mL purified water each. Soil sealing was observed in the first week in 100% LRS and continued to hinder water absorption for the entirety of the growth cycle. Fluid exchange, both liquid, and gas, is fundamental for crop growth as it allows the soil to maintain humidity, exchange gas, and enable root and microbial respiration [2]. On day 16, all seeds had germinated (100 %) (Figure 3). Plants in 50/75/100 % LRS showed signs of xeromorphism: dwarfism, a loss of leaf area, and reduction/lack of shoot branching. These results are consistent with previous experiments using anorthosite and are likely to protect the plant from water loss due to the LRS’s poor water retention capacity [9, 30].

**Figure 3:**
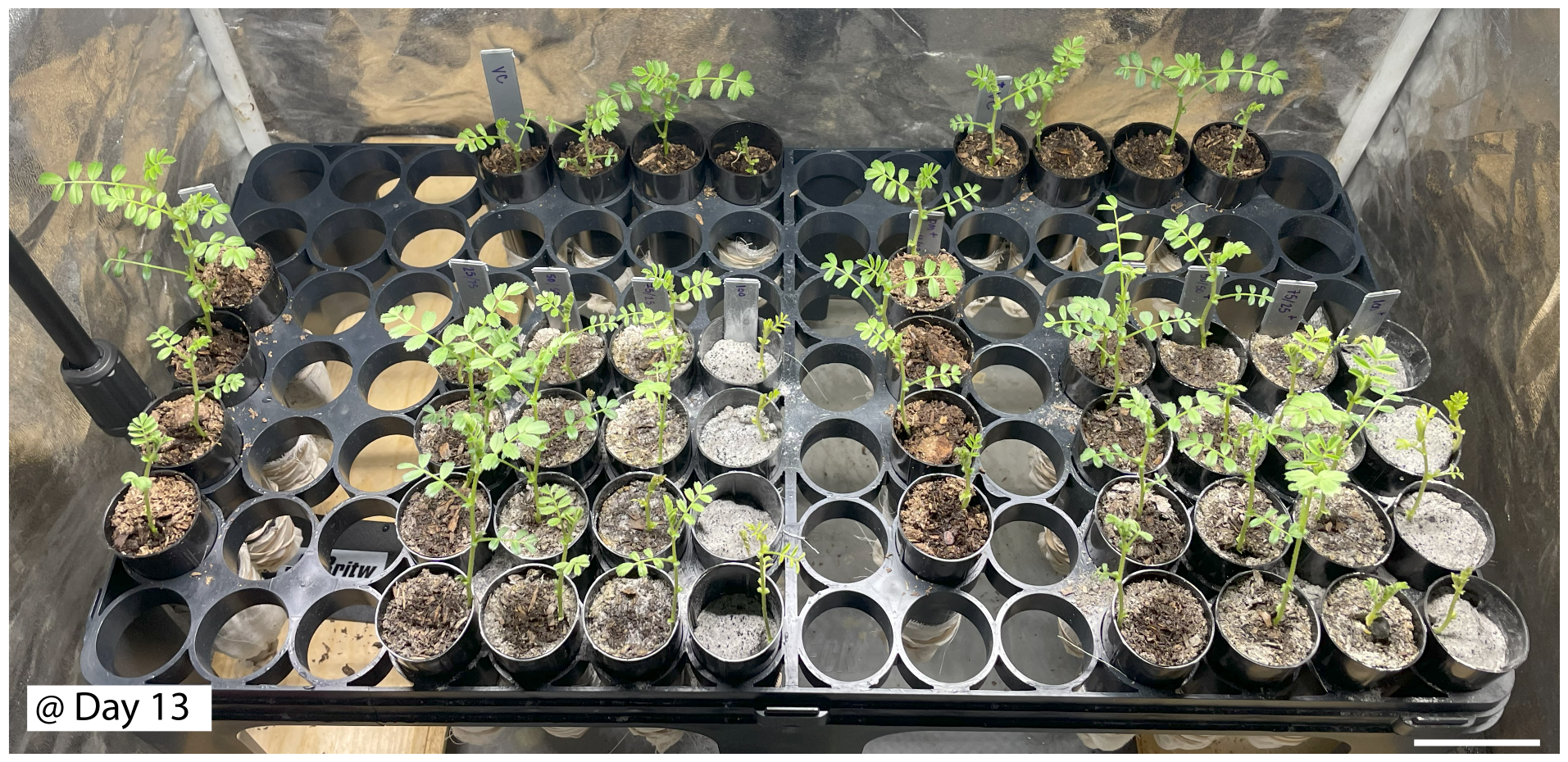
At day 13, there were 100% germination rates. Experiments are labeled as in Figure 2. Potting mix shows large leaves and greater branching, whereas plants in lunar regolith simulant mixtures show signs of xenomorphism, with reduced leaf area, reduced amount of leaf growth, and smaller shoot height. Additionally, chlorophyll levels are lowest in LRS100 (leaf yellowing).

Germination measurements were taken 21 days after 50% emergence in the control group (Figure 4) [29]. These may not account for fungal colonization, as it can take up to eight weeks to see visible plant benefits. Here, we provide a baseline for chickpea seedling emergence in LRS. LRS25, LRS50, and LRS75 compositions had a mixture of small and large particles and showed consistent growth patterns. LRS100 exhibited soil sealing within the first week of planting, limiting water absorption and gas exchange. This group had the highest vertical growth in the germination stage. Studies using *Arabidopsis thaliana* showed that plants that bolt early will senesce early [31]; we hypothesize this rapid growth was a stress response that preceded leaf senescence.

**Figure 4:**
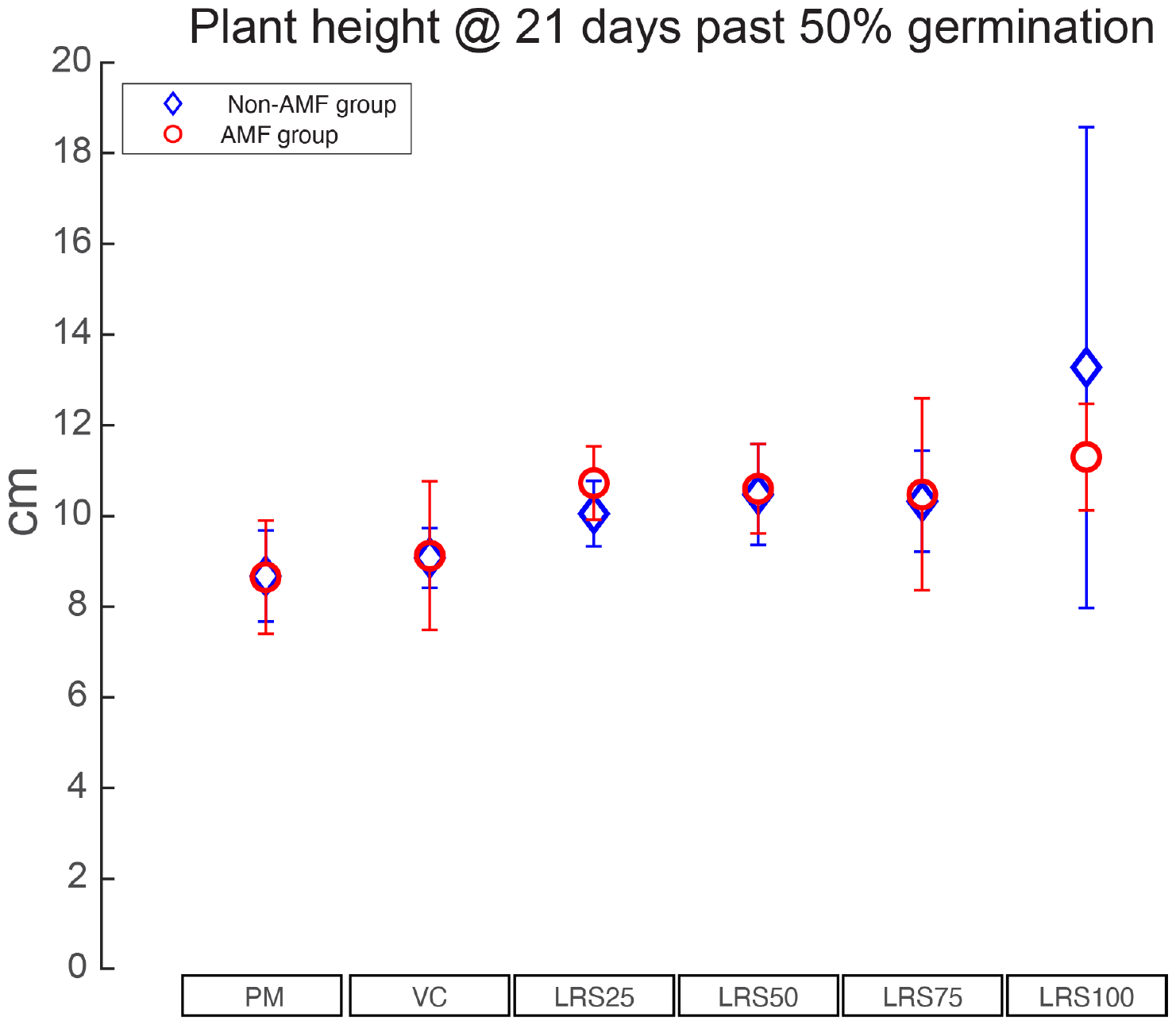
Shoot height at 21 days past 50% germination in the control (PM). All plants germinated (100%) and reached 21 days past 50% germination in the control (PM). Shoot height was measured for all plants (n = 48). Red circles correspond to the AMF-inoculated group; blue diamonds correspond to the non-AMF-inoculated group. Measurements are averaged for each group (n=4), and the standard deviation is represented by error bars on each data point.

Differences in chlorophyll levels were evident through visual observations during the cycle (Figure 5) [32]. Control groups without LRS were used as the baseline, showing that increasing compositions of LRS correlated with heightened signs of stress [32]. By week seven, a visible improvement in chlorophyll levels suggests successful AMF colonization in the inoculated group. At this time, wood dowels were added to support vegetative plant growth. We hypothesize this was necessary due to the small container size in which the plants were growing, limiting root growth, as well as the poor water retention properties of LRS.

**Figure 5:**
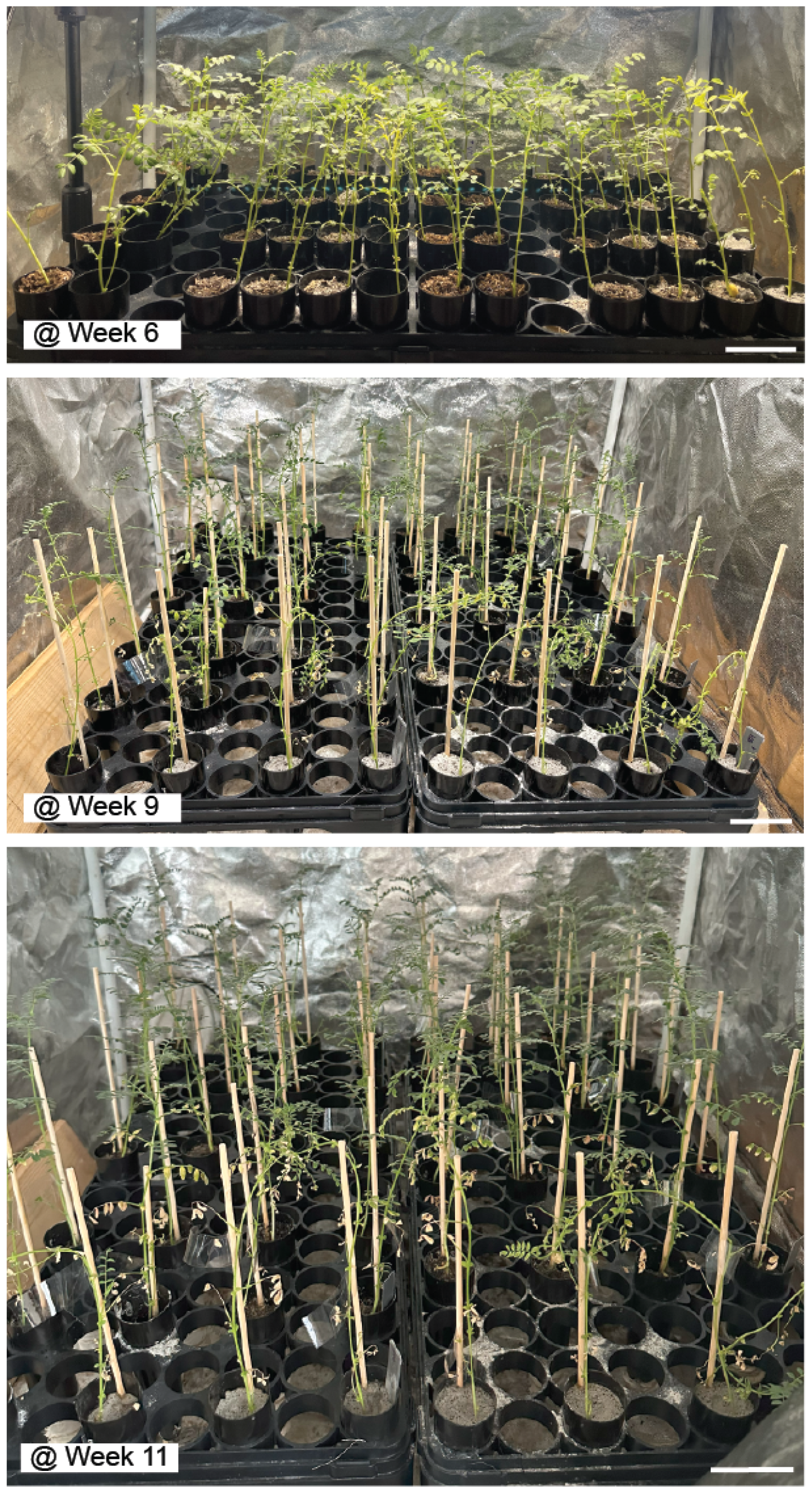
Weeks 6, 9, and 11. Dowels were added in week 7 to add shoot support. Additionally, containers were spaced to allow each plant to grow a canopy. Throughout the cycle (here, showing weeks 3 to 11), we observed different levels of chlorophyll and canopy growth. All plants in LRS100 senesced during vegetative growth. The scale bar represents 5cm.

**Figure 6:**
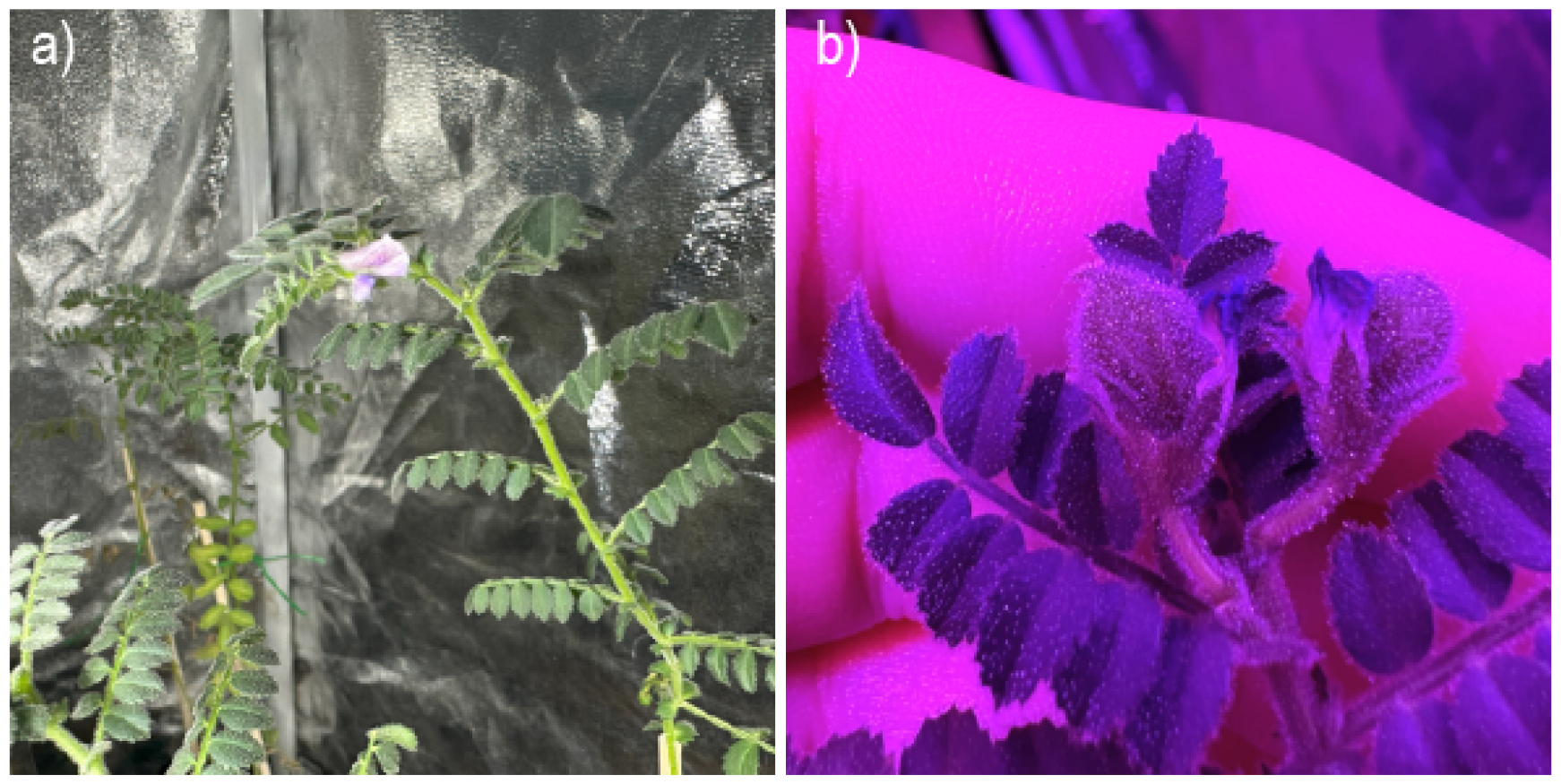
Chickpea flower and seed. Panel a) shows the first flower in AMF-inoculated LRS75, and panel b) shows seeds in AMF-inoculated LRS75.

By week 10, plants from the non-AMF group began to senesce, while inoculated groups began the onset of flowering. Flowering and seed sets were exclusively observed in the AMF-inoculated group 6. All plants in 100 % LRS underwent senescence during the vegetative stage. However, the inoculated group survived an average of two weeks longer. The seed set was delayed, and full maturity took 120 days instead of a typical 100-day cycle. Turgor pressure variations were observed throughout the entirety of the growth cycle. Turgor pressure is an indicator of water deficit and environmental conditions in plants [33]. This was not surprising, considering LRS’s poor hydraulic properties.

## Conclusion

We report the first instance of growing chickpea (*Cicer arietinum*) in lunar regolith simulants. We used soil regeneration techniques common on Earth with LRS for the first time, using both AMF and VC. We also achieved the first documented chickpea yield in an LRS mixture. Our results show that regeneration methods used on Earth soils can help condition lunar regoliths. Despite promising results, all plants in LRS showed signs of chlorophyll deficiency. We assessed the impact of different compositions of LRS on plant growth and development. We achieved flowering in the inoculated 25/75, 50/50, and 75/25 LRS/VC composition. Visual observations of plant stress included stunting, reduced leaf area, limited shoot branching, and death in plants exposed to 100% LRS concentrations.

Throughout the growth cycle, fluctuations in turgor pressure were observed, consistent with the anticipated effects of LR’s poor hydraulic properties. Visual assessments of chlorophyll levels show a correlation with increasing LRS compositions. By week seven, the inoculated group had visible signs of successful arbuscular mycorrhizal fungi (AMF) colonization, suggesting the potential role of AMF in mitigating the stresses induced by LRS. The vegetative stage in 100% LRS resulted in senescence for all plants, but the AMF-inoculated group exhibited a survival advantage, lasting an average of two weeks longer. LRS 50 and LRS 75 inoculated groups demonstrated seed set, revealing successful reproduction under LRS concentrations. These results show the potential of VC amendments and AMF inoculation, in collaboration with the chickpea, to condition LRS and transform it into a productive LRS matrix. Additionally, we showed that AMF can form plant-beneficial associations with plants even in high LRS concentrations, which offers a pathway to reducing heavy metal toxicity in lunar horticultural crops that form relationships with AMF.

This is preliminary data we release to promote open science practices. We will release additional quantitative data that will further contribute to our understanding of plant and microbial responses to lunar regolith simulant encompassing fungal colonization and chemical and physical parameters of the LRS matrix. Additionally, comprehensive data on plant yield will be made available as these experiments are completed. These findings provide valuable insights into adaptive responses, emphasizing the potential role of AMF in enhancing plant resilience in extreme environments.

## Acknowledgements

J.A.A. thanks Dr. Elizabeth Pierson (Texas A&M University) for continued support and mentorship. Dr. Terry Gentry (Texas A&M University) for technical assistance and guidance, and Dr. George Vandemark (Washington State University) for helpful discussions. We thank Jon Howard for support and materials, and Cristy Christie (Black Diamond Vermicompost) for materials and detailed characteristics.

## Author contributions

J.A.A. conceived the experimental idea and designed the experiments. J.A.A performed all experiments. J.A.A. and S.O.S. processed and analyzed the data. J.A.A. and S.O.S. wrote and revised the manuscript.

